# Flexible Analysis of Spatial Transcriptomics Data (FAST): A Deconvolution Approach

**DOI:** 10.1101/2023.05.26.542550

**Authors:** Meng Zhang, Yiwen Liu, Joel Parker, Lingling An, Xiaoxiao Sun

## Abstract

**Motivation:** Spatial transcriptomics is a state-of-art technique that allows researchers to study gene expression patterns in tissues over the spatial domain. As a result of technical limitations, the majority of spatial transcriptomics techniques provide bulk data for each sequencing spot. Consequently, in order to obtain high-resolution spatial transcriptomics data, performing deconvolution becomes essential. Deconvolution enables the determination of the proportions of different cell types along with the corresponding gene expression levels for each cell type within each spot. Most existing deconvolution methods rely on reference data (e.g., single-cell data), which may not be available in real applications. Current reference-free methods encounter limitations due to their dependence on distribution assumptions, reliance on marker genes, or the absence of leveraging histology and spatial information. Consequently, there is a critical demand for the development of highly adaptable, robust, and user-friendly reference-free deconvolution methods capable of unifying or leveraging case-specific information in the analysis of spatial transcriptomics data.

**Results:** We propose a novel reference-free method based on regularized non-negative matrix factorization (NMF), named Flexible Analysis of Spatial Transcriptomics (FAST), that can effectively incorporate gene expression data, spatial coordinates, and histology information into a unified deconvolution framework. Compared to existing methods, FAST imposes fewer distribution assumptions, utilizes the spatial structure information of tissues, and encourages interpretable factorization results. These features enable greater flexibility and accuracy, making FAST an effective tool for deciphering the complex cell-type composition of tissues and advancing our understanding of various biological processes and diseases. Extensive simulation studies have shown that FAST outperforms other existing reference-free methods. In real data applications, FAST is able to uncover the underlying tissue structures and identify the corresponding marker genes.

## Introduction

Spatial transcriptomics has been rapidly expanding during the past decade [1, 2, 3, 4]. It captures gene expression while preserving the spatial structure and information of the tissue. After sequencing, unique coordinates and gene expression levels of each spot are retained. Based on spatial transcriptomics data, we can explore the spatial patterns of expression, tissue architectures, and cell-to-cell interactions [5, 6, 7, 8, 9]. Several techniques of spatial transcriptomics are commonly used. For example, fluorescence imaging-based methods (e.g., merFISH) can provide high-resolution data with gene expression at the almost single-cell level in each spot [10, 11]. However, these methods can only perform sequencing with a limited number of predefined target genes. Next-generation sequencing (NGS) based spatial transcriptomics (e.g., 10X Visium) can provide whole-transcriptome sequencing but with a low-resolution (i.e. 55-100 *µm*) [12, 13, 14]. Throughout this paper, we focus on deconvolution for low-resolution spatial transcriptomics methods. Although deconvolution methods for bulk RNA sequencing (RNA-seq) data have been developed for decades, their generalizations to spatial transcriptomics data are limited due to the difficulties of including the spatial and histology information from the spatial transcriptomics data [15]. New methods explicitly designed for spatial transcriptomics are rapidly emerging [16, 17, 18, 13, 19, 20, 14, 21, 22, 23, 24]. Most of them utilize the reference data that are generated from single-cell RNA-seq data. These reference-based methods offer convincing deconvolution results when the prior knowledge about the reference is accurate, which requires domain knowledge and expertise in biology. Additionally, constructing a reference for deconvolution for a novel problem requires collecting and processing single-cell data when a problem-specific reference is unavailable, making it financially challenging for many labs to get accurate deconvolution results. To overcome the limitations, reference-free methods have been developed. To the best of our knowledge, only a few reference-free methods are available. For example, STdeconvolve is a reference-free spatial deconvolution method built on a latent Dirichlet allocation (LDA) model [23]. STdeconvolve achieves comparable accuracy with reference-based methods and outperforms reference-based methods when golden reference data is not available. LDA encodes the internal distributions for genes across cells and cells over spots. However, a higher drop-out or a smaller number of spots are obstacles for LDA to model such distributions, hence unable to provide highly accurate deconvolution results [23]. As spatial transcriptomics platforms approach single-cell levels, the distributional assumptions placed on the cell types within a spot, may not hold. In addition, the STdeconvolve method mainly relies on the gene expression data of each spot but ignores potential spatial dependencies within the spatial transcriptomics data. The CARD method was initially developed as a reference-based method, but it includes a built-in function CARD-free, which enables deconvolution using only marker genes of cell types [22]. CARD-free can be classified as a semi-reference-based method, because, it utilizes a limited set of marker genes as reference rather than a comprehensive reference dataset. The performance of CARD-free relies on the predefined set of cell types and their corresponding marker genes.

In our study, we propose a novel reference-free approach called Flexible Analysis of Spatial Transcriptomics (FAST), which incorporates gene expression data, spatial data, and histology information to perform deconvolution of spatial transcriptomics data, see Figure 1. We enhance the NMF framework by introducing two penalty terms. The first term incorporates spatial information by utilizing the graph Laplacian matrix, which is constructed by combining spatial and histology data. We introduce a straightforward method to obtain the graph Laplacian matrix in this study. Note that our method is adaptable to any graph Laplacian matrix, allowing for flexibility in its application. The second term imposes a constraint on cell proportions, encouraging their summation equals one. In summary, FAST stands out from existing methods due to its ability to impose fewer distribution assumptions, incorporate spatial tissue structures, and produce interpretable factorization results with greater flexibility. These features make FAST a valuable tool for uncovering the complex cellular composition of tissues and advancing our understanding of various biological processes and diseases that can be elucidated by spatial transcriptomics.

**Fig. 1.**
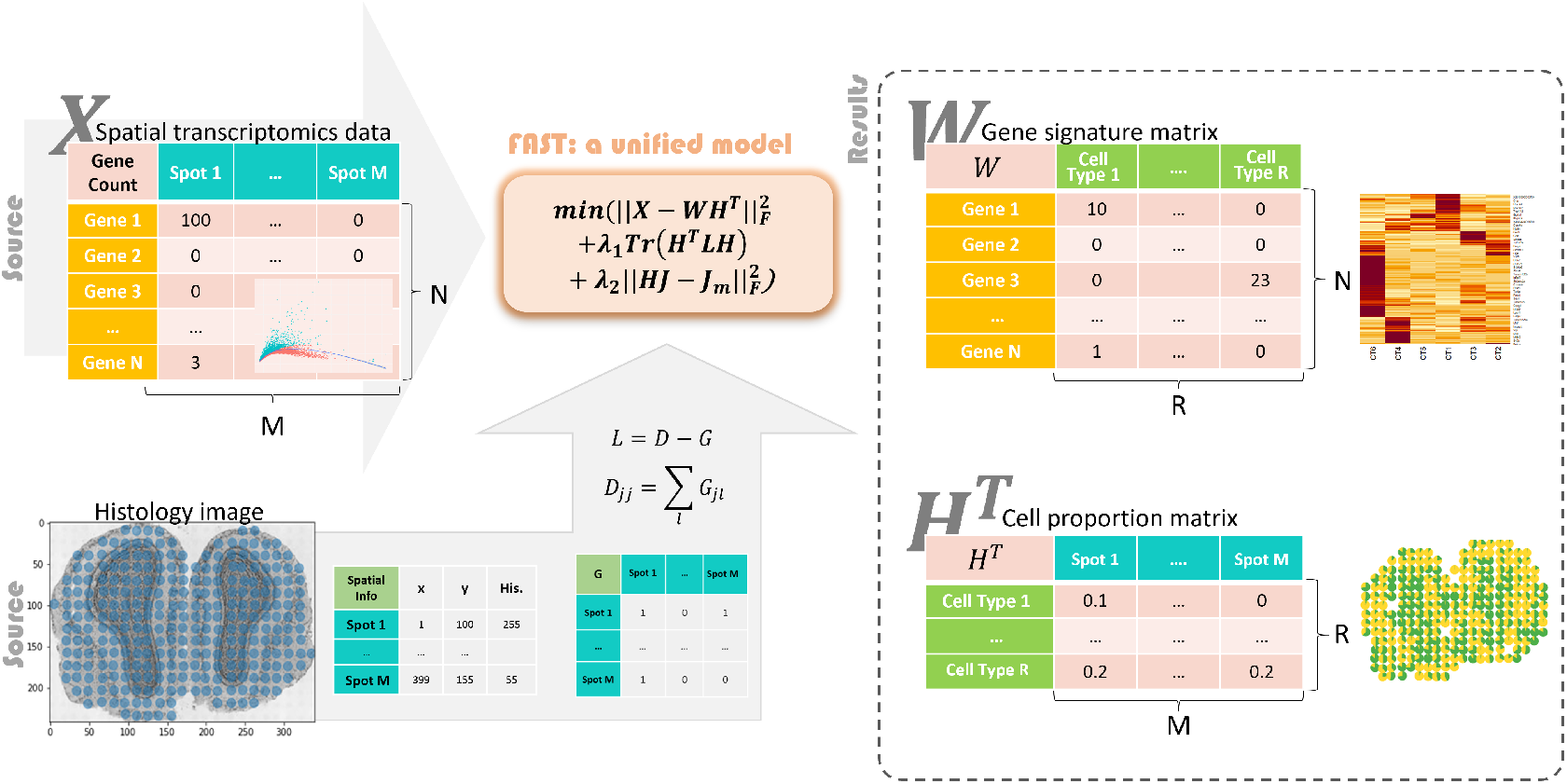
Illustration of the FAST pipeline based on regularized non-negative matrix factorization. X is an N-by-M matrix that contains gene counts per spot, while W and H are a pair of low-rank embeddings of X. W addresses the signature genes of each cell, and H contains the cell proportions in each spot. FAST utilizes X and spatial information to generate X and W in a unified model.

## Methods

FAST is a regional resolute deconvolution method that takes spatial transcriptomics data and a user-defined adjacent matrix as input to produce cell proportions of each spatial spot with the corresponding gene signature matrix as output. The gene expression matrix of spatial transcriptomics data is denoted as an N-by-M matrix *X*_*N×M*_ with N genes as rows and M spots as columns.

Consider the formulation of a simple NMF applied on spatially resolved matrix *X*_*N×M*_,

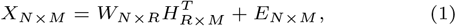

where *W* is an N-by-R matrix that represents the gene signature/transcriptional profile matrix of R cell types, *H* is an M-by-R matrix that represents the abundance of R cell types in M spots, and *E* is the error term. To minimize the error term, the objective function can be expressed as,

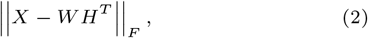

where ||*·*||_*F*_ is the Frobenius norm.

To incorporate the spatial information and the biological nature of the tissues into the objective function in (2), we add two regularization terms and construct the following objective function,

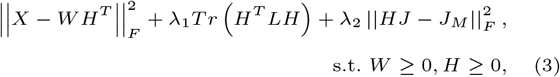

where *Tr* (*H*^*T*^ *LH*) integrates the spatial information of spots with histology information, and 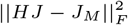 imposes the summation to one penalty of cell proportion estimates for each spot [25]. Their regularized parameters *λ*_1_ and *λ*_2_ control the impact of each term. Particularly, the graph Laplacian matrix is defined as *L* = *D* − *G* where *D*_*jj*_ = ∑_*l*_ *G*_*jl*_, and *G* is the user-defined adjacent matrix for the nearest neighbor networks of spots. *J* is a R-by-M matrix with all elements equal to 1, and *J*_*M*_ is an M-by-M square matrix with all elements equal to 1. The proposed method is flexible in a way that the adjacent matrix can be defined using various approaches. We propose one method in this paper, which is introduced in the next subsection. We solve *W* and *H* in (3) using the updating rules shown in Algorithm 1. Details of the derivation of updating rules can be found in Supplementary Material.

### Algorithm 1

Updating rules for the FAST algorithm

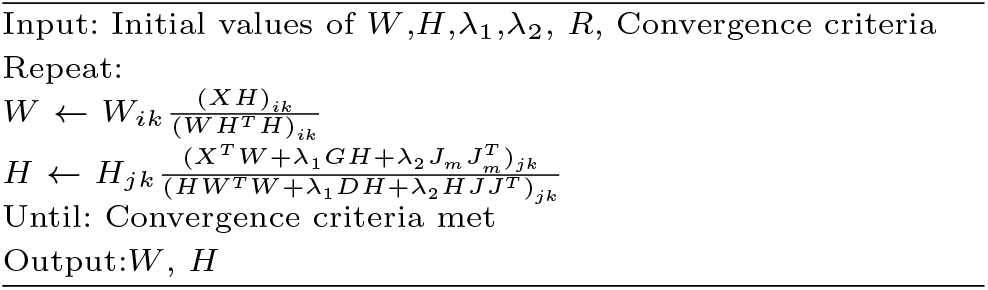

### Construction of the adjacent matrix

The accurate construction of the adjacent matrix is critical for the success of the proposed algorithm. In this paper, we propose a straightforward method that incorporates spatial information and histology data when constructing the adjacent matrix *G*. The adjacent matrix should reflect the local spatial structures of the spots. Intuitively, spots that are physically close to each other are likely to share similar expressed gene sets and cell type distributions. However, this is not always true when organs are biologically segmented into special shapes. For example, blood vessels are tubular structures that can appear elongated or circular when viewed under a microscope. The similarity of spots in the above types of organs cannot be measured solely based on physical distance. We aim to construct an adjacency matrix based on biological proximity rather than physical distance. In order to find a balance between physical distance and intrinsic biology structure, histology images are introduced. They are microscopic images of tissue samples on glass slides stained with various dyes to enhance the visibility of specific features, such as cell nuclei or protein structures.

The proposed method calculates the adjacent matrix by integrating spatial histology and spatial coordinates in Euclidean space. We assume spots that are closer both histologically and spatially tend to have similar cell type distribution. Therefore, we compute Euclidean distances of histology and 2D coordinates of spots. The distance between two spots on histology can be calculated by measuring the difference in their median intensities over a sub-region after converting the images to grey-scale ones. In this work, the sub-region is defined as a 5-by-5 square centered around each spot, and the median intensity of the 25 spots is reserved as the value of the corresponding spot. The entries of the adjacent matrix are given by,

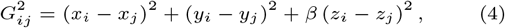

where *x*_*i*_, and *y*_*i*_ are spatial coordinates of the *i*th spot, and *z*_*i*_ is the gray-scaled median intensity of a spot on the histology image. The parameter *β* controls the relative scale of median intensity and spatial coordinates of spots. Some histology images are vague and less informative, and *β* should be assigned with a smaller number in this case. Our recommended *β* is

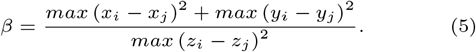

We also use a sparse adjacent matrix to improve the efficiency of the proposed algorithm [25]. Particularly, we only keep the top five largest values in each row of *G*, while the rest of the values are set to zeros.

### Evaluation

Proper annotation of cell types improves the capability of biological interpretation of the results. In the simulation studies, we use a data-driven method to identify the cell type for each factor in *W* and *H*. In particular, we calculate the correlation of each factor in *H* with the true cell proportions of all cell types. The cell type with the highest correlation value is assigned to annotate the factor. The details are shown in Supplementary Material.

To evaluate the performance of the methods in the simulation studies, we utilize multiple evaluation criteria. Average Pearson correlation coefficients were computed to measure the mean correlation between the true and estimated cell proportions over all cell types. Additionally, Root-mean-square error (RMSE) was calculated to measure the differences between the estimated and true cell type proportions.

## Results

We conducted extensive simulation studies and real applications on three spatial transcriptomics datasets to demonstrate the performance and capability of FAST and compared its results with two reference-free methods currently available [22, 23].

### Simulation studies

There exist two popular simulation strategies in generating sptial transcriptomics data [22, 21]. We chose to use the simulation method based on single-cell data. Particularly, we selected cells according to a pre-defined distribution from a single-cell dataset and took the summation of the gene expression levels of the selected cells to fit each spot from the spatial transcriptomics.

We used single-cell RNA-seq data with 18,215 genes and two cell types from the mouse nervous system to construct a spatial transcriptomics dataset on mouse olfactory bulbs with 260 spots [26]. Then, we selected top differentially expressed genes based on the Wilcoxon signed-rank test with an adjusted cutoff p-value 1 *×* 10^−5^, resulting in 5,160 selected genes. A Dirichlet distribution was used to determine the proportions of each selected cell type, see Table 1. We used two cell types of Astrocytes and Neurons. For the 75 spots of the granule cell layer, Astrocytes is the dominant cell type with *α*_1_ = 1, *α*_2_ = 3, and Neurons is the dominant cell type in the 45 spots of the nerve layer. The rest 140 spots from the mitral cell layer have both cell types balanced distributed with *α*_1_ = *α*_2_ = 1.

**Table 1.** Simulation Settings

| Region | Spots<sup>1</sup> | Dominant<sup>2</sup> | Parameters<sup>3</sup> |
| --- | --- | --- | --- |
| Granule cell layer | 75 | Astrocytes | $\alpha_1 = 1, \alpha_2 = 3$ |
| Mitral cell layer | 140 | N/A | $\alpha_1 = 1, \alpha_2 = 1$ |
| Nerve layer | 45 | Neurons | $\alpha_1 = 3, \alpha_2 = 1$ |

<sup>1</sup>Number of spots
<sup>2</sup>Dominant cell type of a region
<sup>3</sup>Parameters of the Dirichlet distribution

The spatially resolved pie chart in Figure 2 (a) shows clear patterns across the three layers of the tissue. Figure 2 (b) shows the scatter plots of the true and calculated proportions of Astrocytes across three cell layers. The closer the dots are to the 45-degree line, the better the performance. Results from FAST are consistently closer to the 45-degree line than the outputs from the other methods. This is further supported by the circular bar charts in Supplementary Material. Figure 2 (c) and (d) show the results of 100 simulation replicates comparisons using Pearson Correlation and RMSE, respectively. FAST demonstrates the highest Pearson Correlation and the lowest RMSE, indicating more accurate performance compared with the other two methods. The average Pearson correlation coefficient of the proposed method was 0.93, with an increase of 0.11 compared with the best result of the other two reference-free methods. The RMSE was 0.15 on average with a corresponding improvement of 0.03. FAST also has the lowest standard deviation (i.e., 0.010 and 0.011) for both measurements, implying consistent and stable performance.

**Fig. 2.**
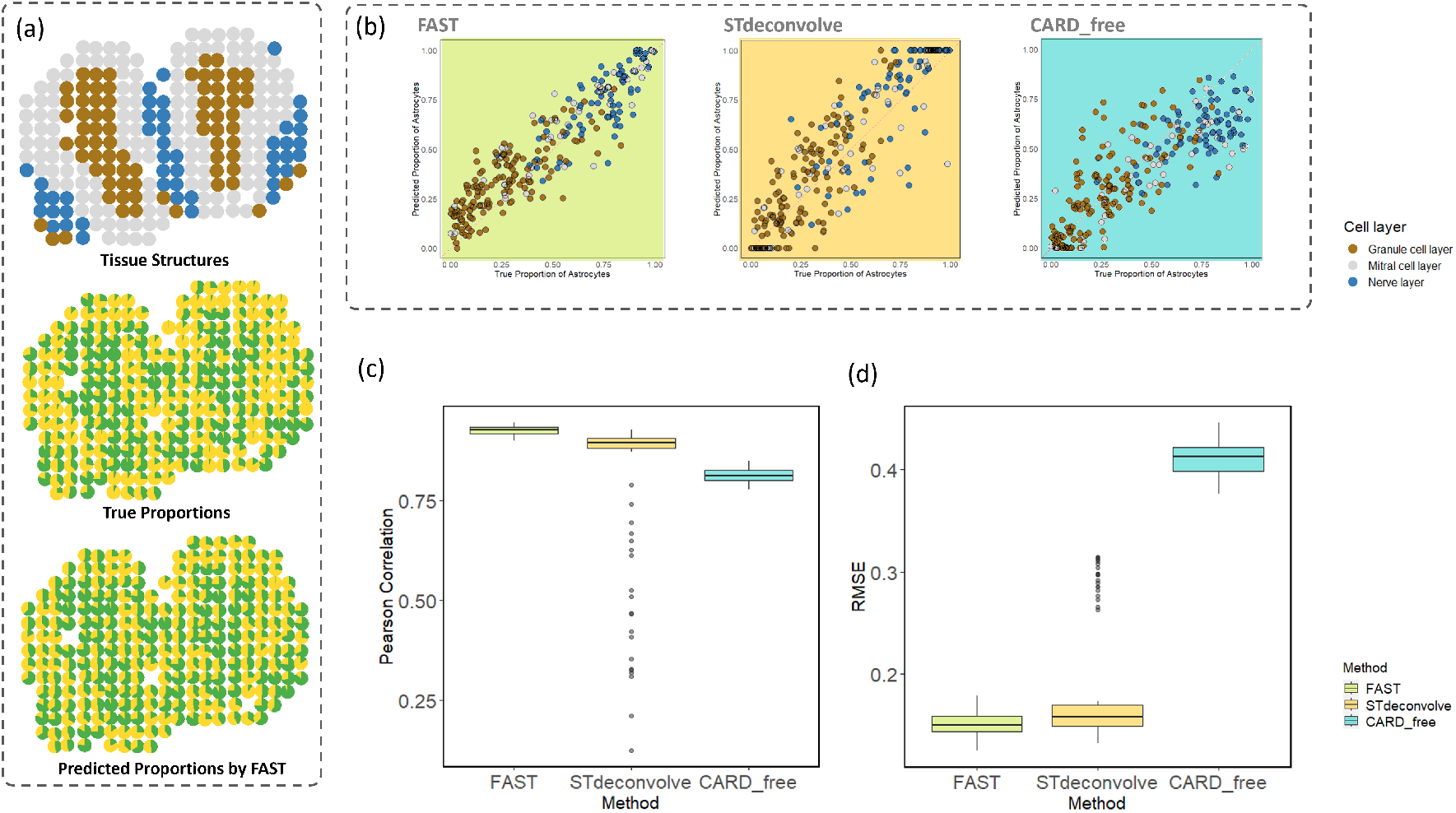
Simulation results. (a) The tissue structures, true proportion pie chart, and predicted proportion pie chart by FAST. (b) Scatter plots showing the distance between predicted and true proportions of the Astrocytes for three methods. (c) Performance comparison of three methods in terms of Pearson correlation for 100 replicates. (d) Performance comparison of three methods in terms of RMSE for 100 replicates.

### Real data applications

We conducted real data analysis for three datasets across two platforms. Two datasets were generated from the spatial transcriptomics platform [27]. The third dataset was generated by the 10X Visium technique with a higher spatial resolution (55 *µ*m).

#### FAST recovers the structures of the mouse olfactory bulbs

The mouse olfactory bulb (MOB) is an important organization of the nervous system located at the front of the brain in mice. It receives and processes signals from olfactory receptor neurons and outputs information to other parts of the system involved in odor detection and processing. Research on MOBs helps researchers to understand the human brain structure and operation of the olfactory system to develop biomimetics smell sensors [28, 29]. The MOB spatial transcriptomics data are well-annotated, which can be served as a good reference when benchmarking MOB spatial transcriptomics analysis.

Although true proportions of cell types in each spot are not available in this dataset, we can still use the annotation of MOB layers as a reliable reference for performance evaluation. There are twelve replicates for this data. Since the downstream analysis based on each replicate achieves very similar results [27], we only selected one replicate (i.e., replicate eight) for data analysis. We used a build-in function in the R package Seurat to select highly variable genes across spots [30]. Five thousand spatially variable genes were selected out of 16,218 genes. We chose the top five nearest neighbors to obtain a sparse adjacent matrix in FAST. MOB is structured in layers with discriminable cell types and functions. In this tissue slide, five layers are annotated from the outermost layer inward as the olfactory nerve layer (ONL), the glomerular layer (GL), the outer plexiform layer (EPL), the mitral cell layer (MCL) and the granule cell layer (GCL). Figure 3 (a) shows the annotations of different layers for 260 spots. Figure 3 (b) shows the clustering results using the K-means clustering algorithm based on cell proportion matrix *H* from the FAST algorithm. The heatmap of cell proportion matrix *H* is shown in Figure 3 (c), in which different layers are well separated based on the dominant cell types. For instance, the first cell type (CT1) was the dominant cell type of the olfactory nerve layer, which was illustrated in Figure 3 (e). To demonstrate the capability of FAST in detecting marker genes, we generated a heatmap of gene expression profiles of all cell types, as shown in Figure 3 (d). The distinct and coherent grouping of genes observed in the heatmap demonstrates the biologically interpretable results obtained from FAST. Our algorithm can also identify marker genes. In Figure 3 (f), the marker gene Kctd12 of CT1 was only expressed in the spots associated with the olfactory layer [27]. The visualization provides evidence that FAST can recover the heterogeneity of tissue structures of MOB at the cell and gene expression levels. We present additional comprehensive gene and cell type coexpression plots in Figure 4 [22, 27, 31]. Patterns of gene expressions are visualized together with the dominant cell types across spots which are represented by dot size. For example, in the last panel of Figure 4, the heatmap visualizes the expression pattern of gene Penk which has higher gene expression levels in dominant CT3. Based on this observation, we can conclude that Penk serves as a marker gene for CT3.

**Fig. 3.**
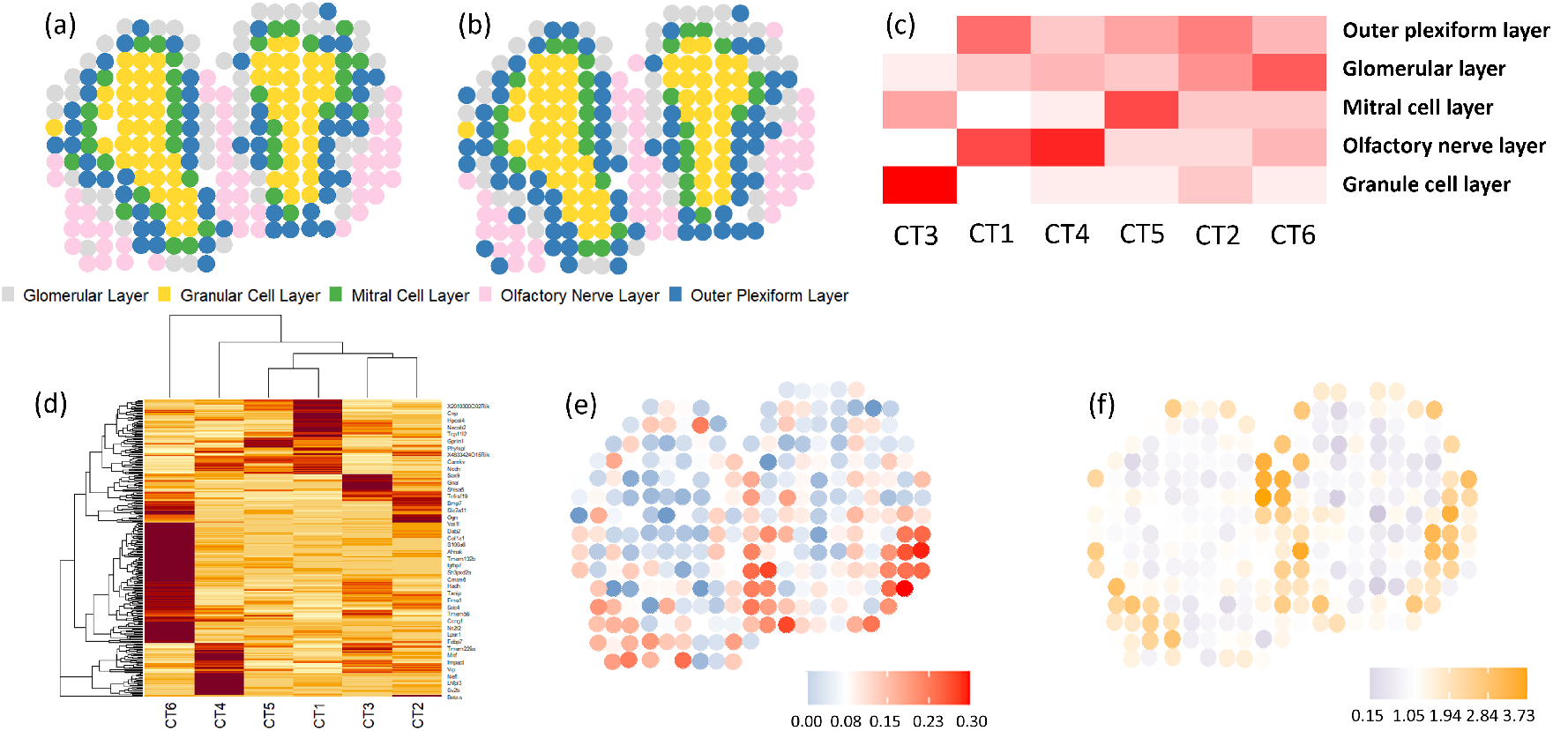
Results of FAST for the MOB data. (a) Annotated layers of MOB. (b) Clustering results using the cell proportion matrix output by FAST. (c) Heatmap of the factor matrix *H*. (d) Heatmap of the gene factor matrix W. (e) Scatter plot of the proportions of cell type 1 (CT1). (f) Spatial scatter plot of the expression levels of gene Kctd12.

**Fig. 4.**
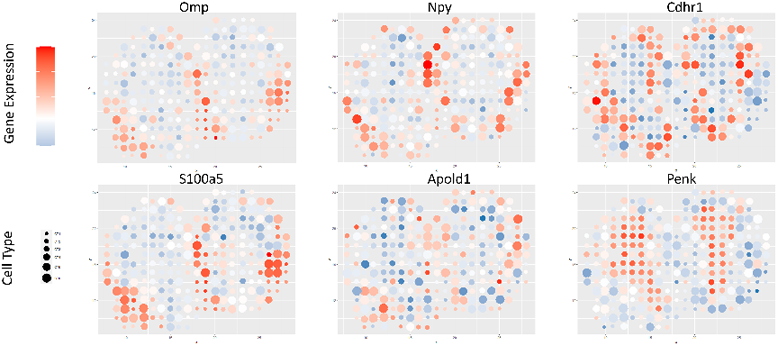
Six marker genes are shown across six factorized cell types in MOB data analysis. The color bar represents the gene expression level, and the dot size indicates the estimated proportion of each of the six cell types by FAST.

#### FAST distinct cancer regions in different stages

The second study provides downstream analysis based on the deconvolution of human breast cancer tissues aiming to assist cancer diagnosis and treatments using spatially resolved transcriptomics data, see Figure 5 (a) [27]. As the most common cancer type, breast cancer has the largest incidence rate in women worldwide [32]. Identifying cellular heterogeneity greatly assists cancer diagnosis [33]. Ductal carcinoma in situ (DCIS) is a non-invasive breast cancer commonly confined to the milk ducts, and invasive ductal carcinoma (IDC) is invasive and can spread to other body parts. Distinguishing between the two types of breast cancer is critical for determining the best treatment from all the options like surgery, radiation therapy, and chemotherapy [34, 35]. In literature, partial annotation is available for DCIS, IDC, and non-malignant regions, see Figure 5 (c) [36]. K-means clustering based on the estimated cell proportions of FAST can recover the annotated spots and extend the annotations to those areas that were previously unclear, see Figure 5 (b) and (d). The cell abundance analysis showed the dominant cell types in different regions, see Figure 5 (e). For instance, cell type 5 (CT5) was the only cell type with high abundance in both DCIS and IDC clusters. In addition, CT1 and CT10 were two of the signature cell types in the DCIS cluster, while CT2 and CT4 were the signature cell types of the IDC cluster. We also conducted a gene enrichment analysis on dominant cell types of tissue clusters [37, 38]. Figure 5 (g) are pathways of the common and distinct cell compositions between the DCIS and IDC clusters. The pathways enriched in CT5 exhibit a high degree of consistency with existing literature on breast cancer pathways (e.g., ECM-receptor interaction pathway), providing further evidence of the biological relevance of this cell type in the context of breast cancer [39]. In addition, several studies have indicated a potential association between the PI3K-Akt signaling pathway and breast cancer progression. We observed a stronger activation of the PI3K-Akt signaling pathway in CT4 compared to other cell types. CT4, identified as a discriminant cell type in the IDC and DCID regions by FAST, provides new evidence of the distinguishing power of this signaling pathway in breast cancer. A list of the presented pathways can be found in Supplementary Materials.

**Fig. 5.**
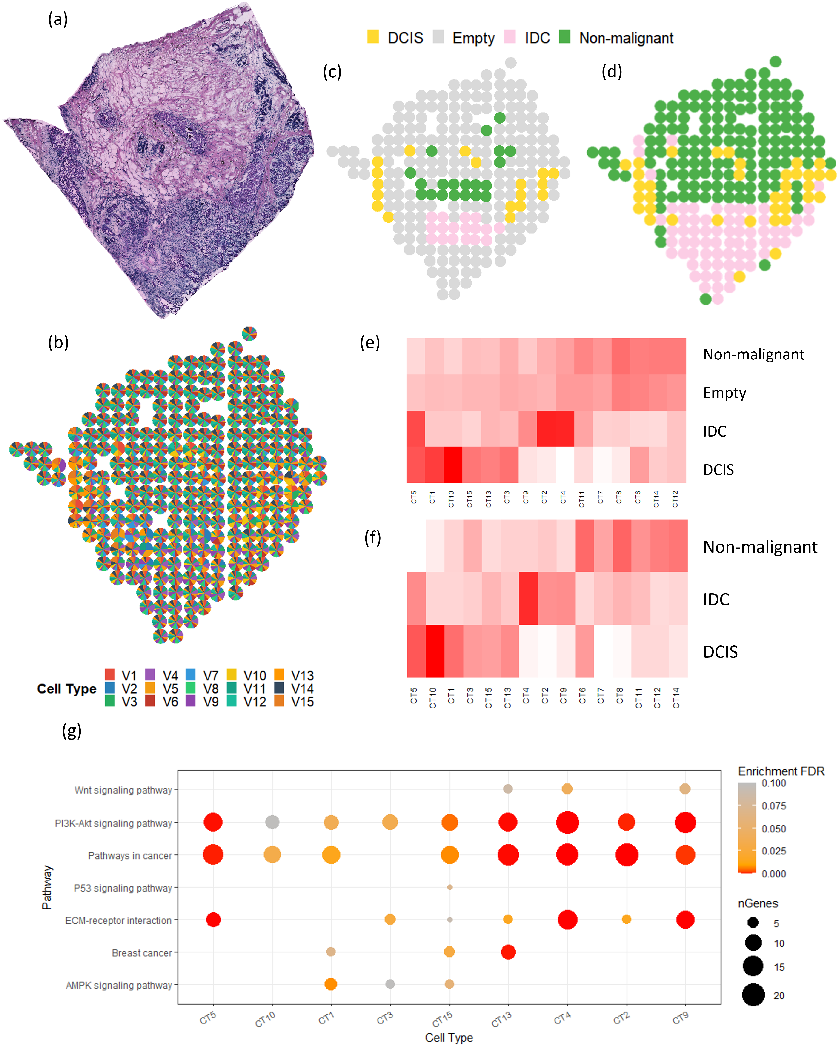
Results of FAST fot the breast cancer data. (a) Histology image of the breast cancer tissue slide. (b) Cell proportion pie chart by FAST. (c) Annotated tissue types. (d) Tissue types generated by FAST. (e) Cell type by annotated tissue region heatmap. (f) Cell type by FAST tissue region heatmap. (g) Dotplot of gene enrichment results.

#### FAST can be applied to higher resolution data to recognize known brain structures

FAST can also be efficiently applied to transcriptomics data with higher spatial resolution. We analyzed transcriptomics data of a coronal section from a mouse sequenced by 10X Visium technology with 2,702 spots and 32,285 genes (Figure 6 (a)). We set the number of cell types to 20 and conducted deconvolution using FAST. Figure 6 (b) is the pie chart showing the proportions of 20 cell types. To enhance the visualization of a cell distribution across all spots, we generated a proportion map for each cell type individually. This allowed us to observe the relative abundance and compare the distribution of a specific cell type with the tissue type classified by Allen Brain Atlas [40]. Figure 6 (c) and (d) show the spatial distributions of CT2 and CT3, respectively, which map to the hypothalamus and Isocortex of the mouse brain. Hypothalamus is located near the base of the mouse brain that is related to many physiological processes like hunger, thirst, etc. Isocortex, often referred to as neocortex, is located on the surface of the brain and controls higher cognitive functions such as perception and language.

**Fig. 6.**
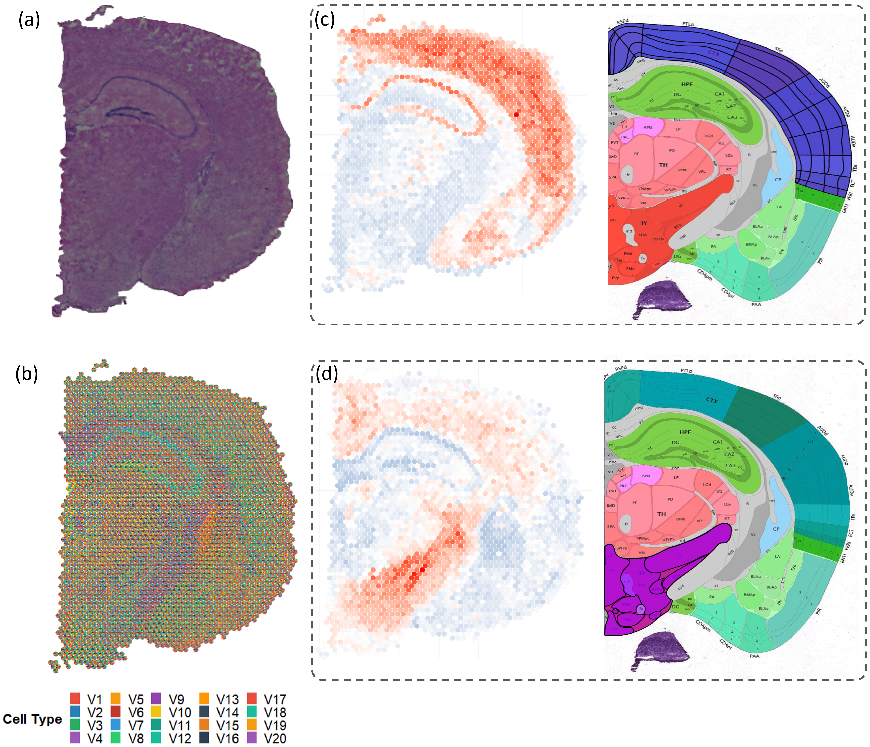
Results of FAST for the mouse brain data. (a) Histology of the mouse brain tissue slide. (b) Cell proportion pie chart by FAST. (c) The scatter plot of the proportion of CT2 and hypothalamus of the mouse brain. (d) The scatter plot of the proportion of CT3 and Isocortex of the mouse brain.

## Discussion

In this article, we developed FAST, a novel reference-free deconvolution method for spatial transcriptomics data based on regularized NMF that integrates gene expression levels, spatial tissue structures, and histology patterns into one unified NMF model. The spatial and histology data are incorporated into the model through a graph regularization term, which utilizes a user-defined adjacent matrix. We further introduced an additional penalty on the proportion matrix to encourage the appropriate scale and uniqueness of both factorized matrices for the first time. FAST surpasses other reference-free deconvolution methods in terms of estimating cell proportions in the simulation study and showcases its potential to unlock new insights and opportunities for in-depth biological research in real data applications.

To enhance the capabilities and applicabilities of FAST, there are several directions that can be explored for future extensions and improvements. First, improving the adjacent matrix with extra information. The current adjacent matrix is calculated using spatial coordinates and the intensities of histology. A promising direction for improvement lies in defining the similarity of two spots using deep learning feature (i.e., texture) detection. Color alone is not the sole resource that can be extracted from an image, and it is vital to incorporate a comprehensive observation of histology. In addition, users have the flexibility to modify the adjacent matrix using their domain knowledge of the tissue structure and control the impact of the graph regularization term according to the level of information that the adjacent matrix contains. Second, the current updating rules are derived using the Frobenius norm in the formulations, a straightforward improvement would be to replace the Frobenious norm with Kullback-Leibler divergence and compare the performance with the current framework [41]. Last, FAST is not limited to a specific domain, and it can be applied to other deconvolution applications with minor modifications on the adjacent matrix. For example, FAST could easily be extended to any problem requiring proportional devolution types of tasks. This will allow users to benefit from the improved stabilization of the NMF algorithms by inducing a sum-to-one penalty term.

## Author contributions statement

M.Z. and X.S. conceived the idea, M.Z. analyzed the results. All authors wrote and reviewed the manuscript.

